# Accurate *in silico* confirmation of rare copy number variant calls from exome sequencing data using transfer learning

**DOI:** 10.1101/2022.03.09.483665

**Authors:** Renjie Tan, Yufeng Shen

**Affiliations:** Department of Systems Biology, Columbia University, New York, NY, USA; Department of Biomedical Informatics, Columbia University, New York, NY, USA; JP Sulzberger Columbia Genome Center, Columbia University, New York, NY, USA

**Author notes:** Correspondence: R.T. and Y.S.

## Abstract

Exome sequencing has been widely used in genetic studies of human diseases and clinical genetic diagnosis. Accurate detection of copy number variants (CNVs) is important to fully utilize exome sequencing data. However, due to the nature of noisy data, none of the existing methods can achieve high precision and high recall rate at the same time. A common practice is to perform filtration with quality metrics followed by manual inspection of read depth of candidate CNV regions. This approach does not scale in large studies. To address this issue, we present a deep transfer learning method, CNV-espresso, for confirming rare CNVs from exome sequencing data *in silico*. CNV-espresso encodes candidate CNV regions from exome sequencing data as images and uses convolutional neural networks to classify the image into different copy numbers. We trained and evaluated CNV-espresso on a large-scale offspring-parents trio exome sequencing dataset, using inherited CNVs in probands as positives and CNVs with mendelian errors as negatives. We further tested the performance using samples that have both exome and whole genome sequencing (WGS) data. Assuming the CNVs detected from WGS data as proxy of ground truth, CNV-espresso significantly improves precision while keeping recall almost intact, especially for CNVs that span small number of exons in exome data. We conclude that CNV-espresso is an effective method to replace most of manual inspection of CNVs in large-scale exome sequencing studies.

## Introduction

Copy number variation refers to >50 bp deletion and duplication in the human genome (1,2). Copy number variants (CNVs), especially for rare CNVs, have been implicated in human diseases and phenotypic diversity (3–7). CNVs can be identified by many genomic technologies such as fluorescent in situ hybridization (FISH), array comparative genomic hybridization (CGH), single-nucleotide polymorphism (SNP) array, next-generation sequencing (NGS), and long-reads sequencing technologies (1,2,8,9). Among NGS technology, exome sequencing technology only performs sequencing on the coding regions which account for ∼2% of human sequence. Exome sequencing has the advantages of high efficiency, low cost, and less storage spaces compared with whole genome sequencing (WGS) technology. Therefore, exome sequencing is widely used as the main sequencing approach by numerous genomic studies of human diseases (10–12). Recently, many exome-sequencing-based CNV detection methods have been developed (13–23). However, the accuracy of CNV detection from exome sequencing data is challenging, especially for small CNVs. The number of CNVs predicted by different methods varied from several to hundred CNVs per sample in which many of them are inconsistent calls (24,25).

To achieve a better performance, a common approach is to call CNVs from exome data using one or multiple methods, followed by empirical filtering based on summary metrics from these methods and then manual visualization of read pileups in regions that harbor candidate CNVs. This approach does not scale in studies with large sample size where manual inspection of large number of candidate CNVs can be extremely time-consuming (26–28). Furthermore, the quality of the manual inspection method is dependent on the experience of investigators, but even experienced investigators may make inconsistent judgement in different settings or time.

Here we describe a new method, CNV-espresso, that can perform *in silico* confirmation of candidate rare CNVs by the same read depth summary figures used for manual visualization. The core model of CNV-espresso is a deep convolutional neural network (CNN). We represent a candidate CNV by an image showing the normalized read depth of the carrier sample and its corresponding reference samples. CNV-espresso uses CNNs pre-trained by a large amount of image data (29) to perform transfer learning for predicting copy numbers as a classification question. In this study, we used a large-scale family-based exome sequencing dataset to create training data and evaluate the performance on samples with both exome and whole genome sequencing data. We investigated whether CNV-espresso can improve accuracy beyond common empirical filters and how the improvement depends on the size and type of CNVs.

## Results

### Overview

**Figure 1** shows the overall workflow of this study. We construct training data by leveraging Mendelian inheritance of offspring-parents trio exome sequencing data. The core of CNV-espresso is a deep convolutional neural network model. We perform transfer learning on a pre-trained image recognition model (29) using the constructed CNV, and then evaluate the performance of CNV-espresso on exome samples with WGS data as proxy of ground truth and an experimentally validated dataset.

**Figure 1.**
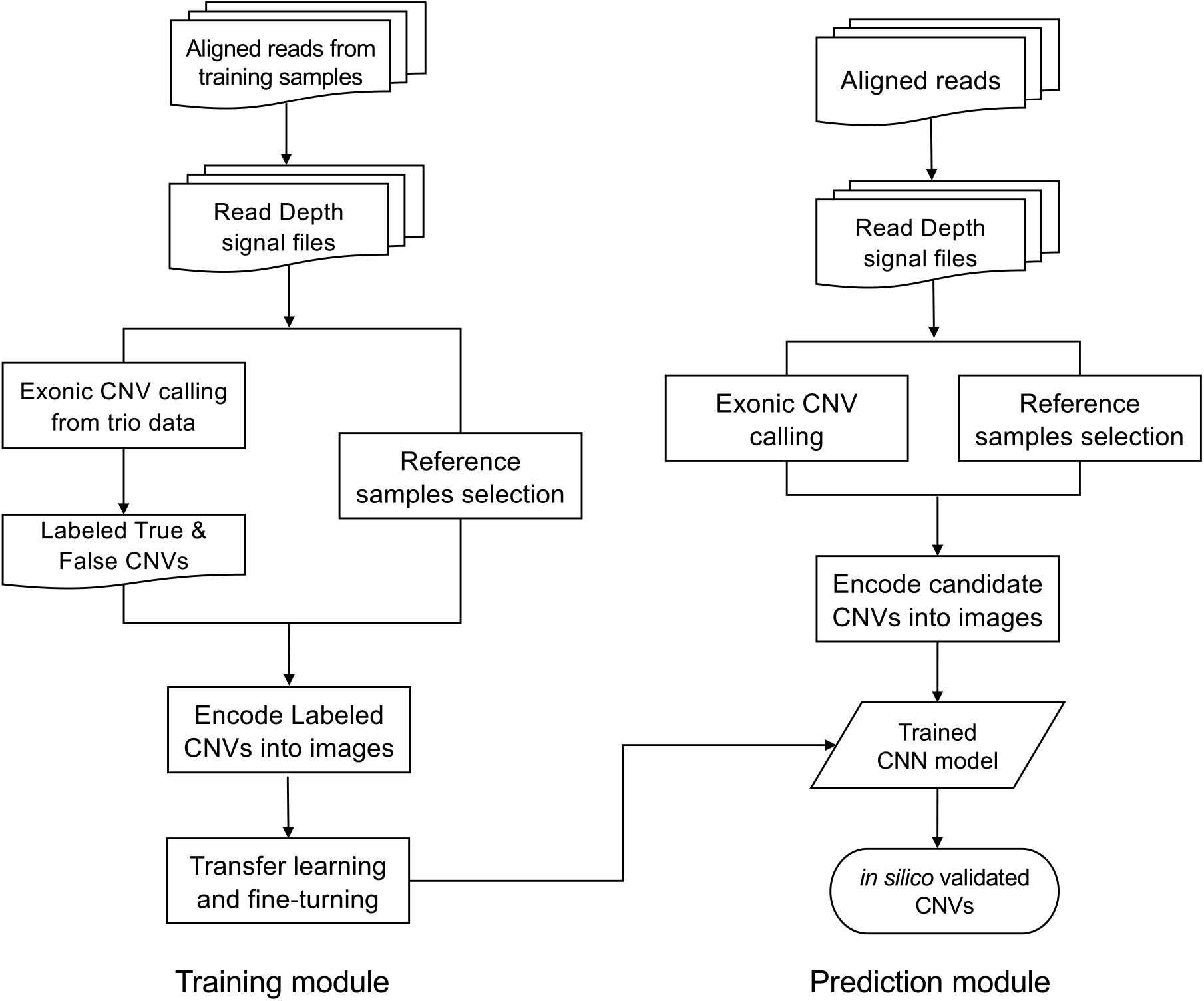
The workflow of CNV-espresso.

### Training data

We obtained 27,270 exome sequencing samples from a study on autism spectrum disorder cohort (Simons Foundation Powering Autism Research for Knowledge, SPARK) (12). We used XHMM (13), CANOES (16), and CLAMMS (19), three exome CNV callers that have different statistical models, to call candidate CNVs. To obtain training data without large-scale confirmation experiments, we used the Mendelian rule of inheritance to construct a high-confidence CNV call set. Specifically, in each family, we assume that most of CNVs called in both offspring and at least one parent that pass baseline quality filters (See Methods) are true CNVs that can be used as *Positives* in training. CNVs called only in offspring but not in either parent are Mendelian errors. These Mendelian error calls include false positives in offspring, false negatives in parents, and true *de novo* CNVs. To refine the Mendelian error calls, we use baseline quality metrics (See Methods) to remove false negatives in parents. For instance, we required ‘NQ’ in both parents to be greater than a stringent threshold to achieve high probabilities of no CNVs in these regions. Meanwhile, based on recent studies, the real *de novo* CNV rate in the human genome is very low (30–32). To minimize the chance of true *de novo* CNVs in the training data, we further excluded candidate CNVs identified by multiple callers, since we found that CNVs identified by multiple callers were likely to be real (**Supplementary Figure 1**). Then we assume that most of the remaining Mendelian errors are mainly composed of false positives in offspring and we use these refined Mendelian error calls as *Negatives* for training. We listed the detailed filtering approaches, quality scores and their thresholds used in this process in **Supplementary table 1**. CNV-espresso is designed to confirm rare CNVs.

Therefore, we excluded CNVs with frequency over 1% in the cohort. As candidate CNVs came from three callers, we first grouped the overlapping candidate by CNV types and genomic coordinates, then resolved the breakpoint conflicts by analyzing the read depth ratios between inside of CNV regions and CNV boundary regions for all possible breakpoints. We processed the merging steps by an external tool (25) and merged different source of CNV calls into an unified call set.

In all, we obtained a training data set with 22,008 CNV calls, including 15,534 positives and 6,474 negatives (**Figure 2**). We randomly partition the data into training (80%) and testing (20%).

**Figure 2.**
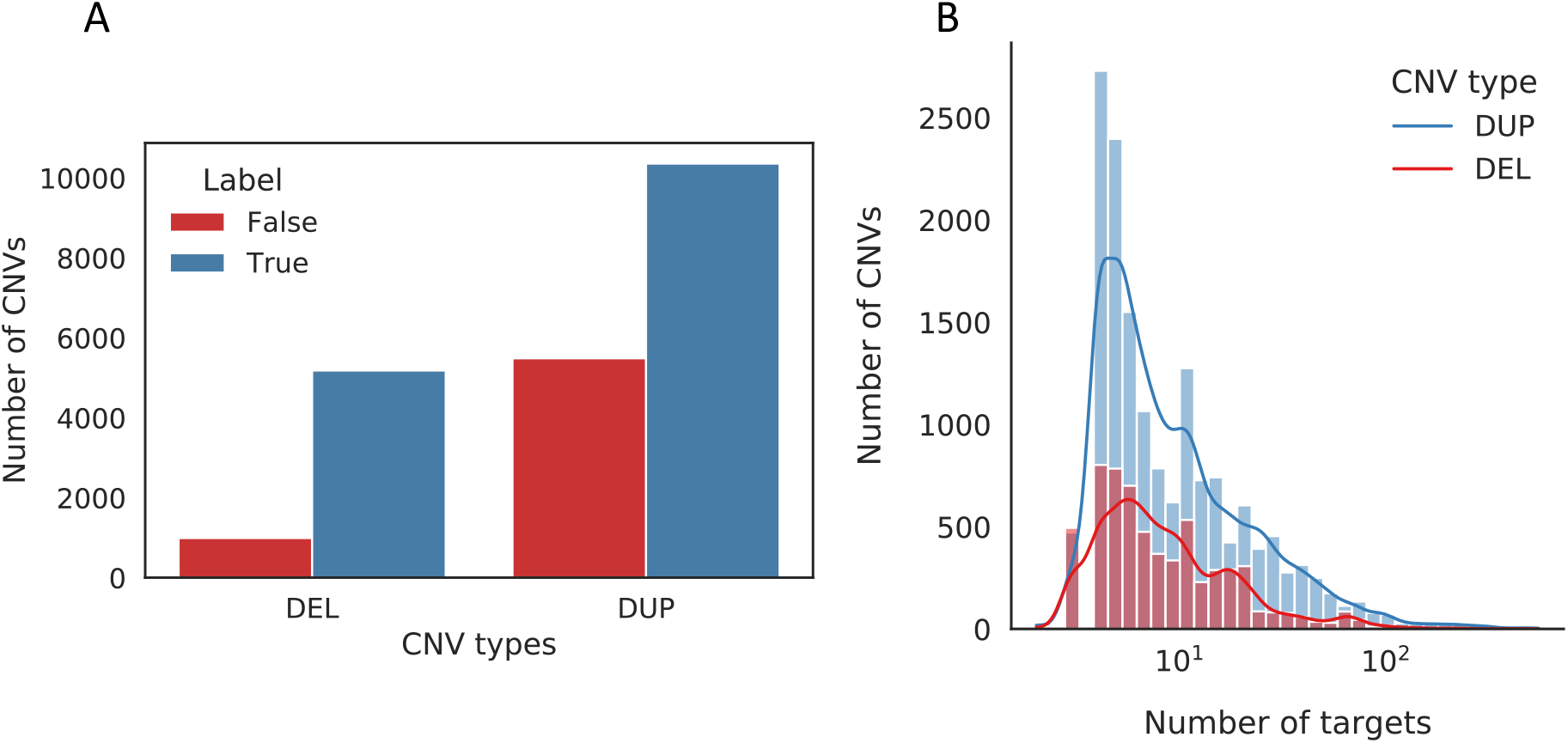
The distribution of CNVs from SPARK dataset. (A) The distribution of CNV types with different labels. (B) The distribution of CNVs with different number of targets. We shuffle the dataset and split CNVs into random training (60%), validation (20%) and testing (20%) data.

### Model performance

We leveraged the transfer learning approach, taking pre-trained models learned from computer vision datasets on our exonic CNV confirmation task. We considered three pre-trained CNN models: a generic CNN model, MobileNet v1 (29), and ResNet50 (33). The generic CNN model includes six convolutional layers, and it can achieve 0.84 accuracies on the CIFAR-10 dataset (34). All other models have been applied to genomics recently (26), (35). Overall, all three models achieved acceptable classification performance. Among them, MobileNet v1 achieved the highest performance with minimum parameters and shortest model training time (**Table 1**).

**Table 1.** Performance comparison and details of three CNN models. We calculated the mean accuracy, F1-score, and GPU time in training data with 5-fold cross-validation.

| Model | Depth | #Parameters | Accuracy | F1-score | GPU time (Minutes) |
| --- | --- | --- | --- | --- | --- |
| A generic CNN | 16 | 22,292,643 | 0.87 | 0.87 | 10 |
| MobileNet v1 | 90 | 3,231,939 | 0.90 | 0.90 | 9 |
| ResNet50 | 178 | 23,593,859 | 0.76 | 0.76 | 15 |

Therefore, we selected MobileNet v1 as our transfer learning base model in the following analysis.

After the transfer learning and fine-tuning steps (See Methods), we evaluated the performance of the refined model in the testing data. The overall F1-score is 0.92, and the model achieves an area under the curve (AUC) of the receiver operating characteristic (ROC) curves at 0.99 (**Figure 3A**). Confusion matrix shown that our model can successfully classify most of deletion, duplication, and diploid predictions (**Figure 3B**). We further evaluated our model on CNVs with the different number of target categories and different CNV types separately. Results shown that our model can achieve good and stable performances on confirming CNVs among different number of target categories (**Figure 4**). Meanwhile, our model also achieved good performances on both deletion and duplication respectively (**Supplementary figure 2**).

**Figure 3.**
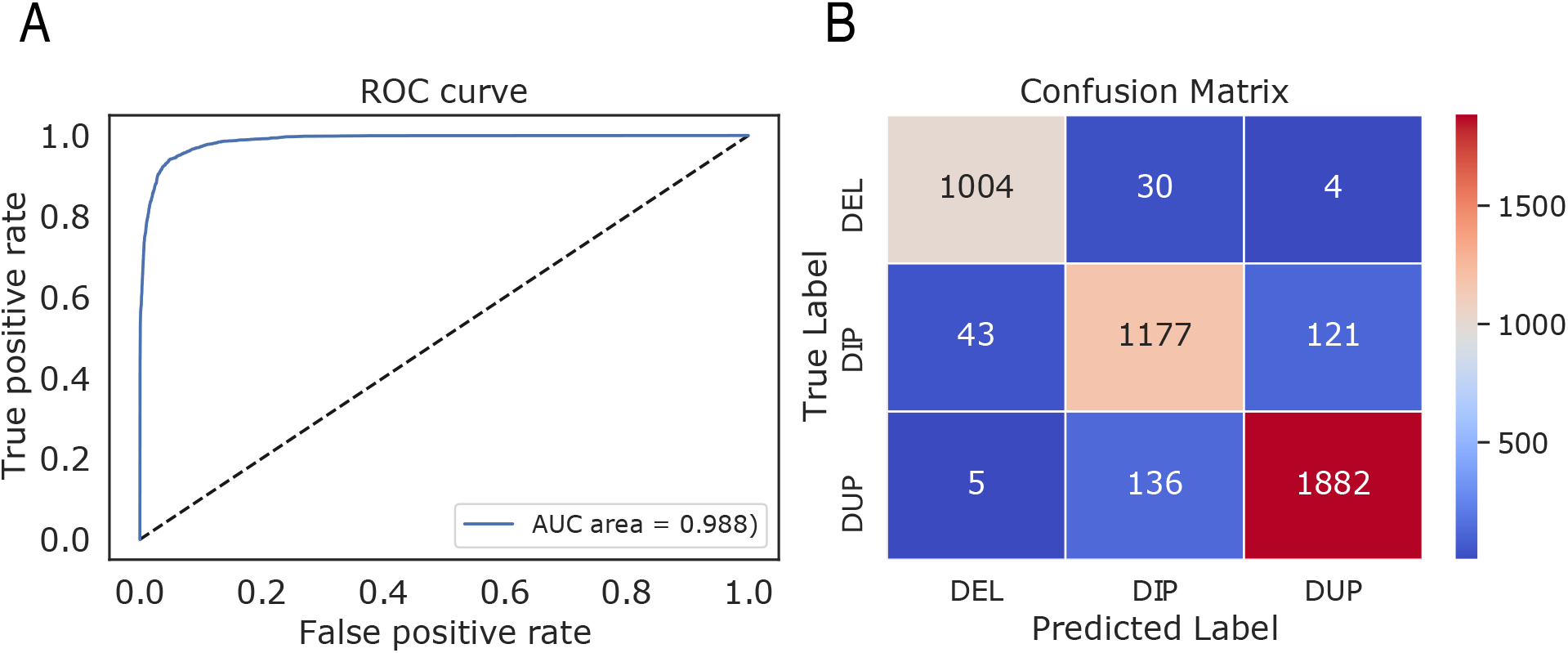
Model performance on SPARK dataset. (A) ROC curve of MobileNet after transfer learning and fine-tuning steps. (B) Confusion matrix.

**Figure 4.**
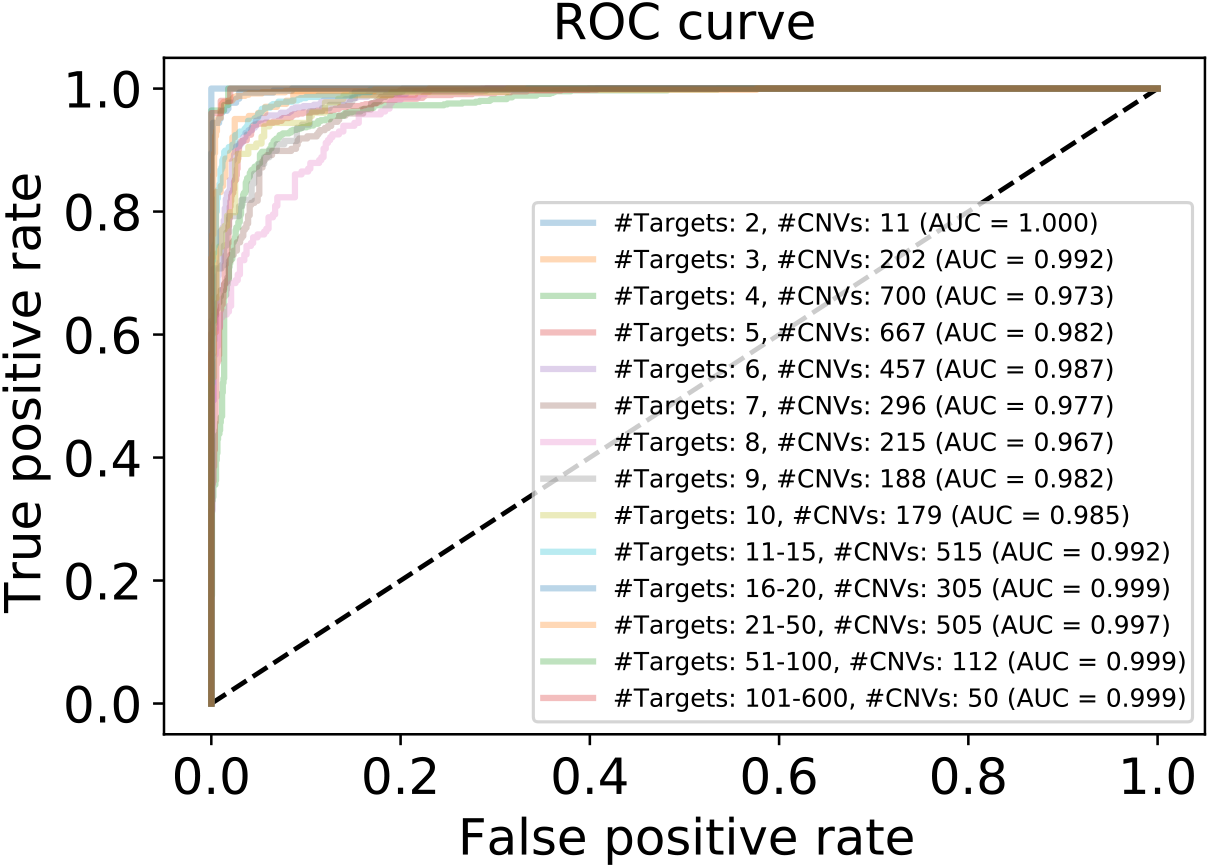
ROC curves for different number of targets.

### Performance of *in silico* confirmation of CNVs

Whole genome sequencing data has effectively complete coverage of genomic regions and more even coverage than exome sequencing data. Therefore, high-confidence CNVs identified from whole genome sequencing data of the same individuals can be used as an approximate gold standard to assess the accuracy of CNVs called from exome data. In this study, we assessed the performance of CNV-espresso on 569 individuals with both exome sequencing and whole genome sequencing data. For these 569 individuals, we used XHMM (13) to call putative CNVs from exome sequencing data, and used Canvas (36) and Manta (37) to identify CNVs from whole genome sequencing data (see Methods). Given that Canvas and Manta are two complementary CNV callers in terms of different input signals and statistical models, we took the union of the high-confidence CNVs identified by the two callers as the standard dataset. We treated CNV predictions from exome sequencing data as true positives if at least 50% of the prediction overlapped with corresponding ones in the standard dataset. We treated the calls without overlap as false positives.

To assess the *in silico* confirmation performance, we classified XHMM-called CNVs using CNV-espresso, and then calculated the precision and recall of resulting CNVs in bins grouped by CNV sizes (number of spanned exons). We considered XHMM-called CNVs at variable quality threshold (SQ estimated by XHMM) (see Methods). Results showed that CNV-espresso can improve the precision with every SQ threshold among different CNV sizes (**Figure 5 A, B, C**). Particularly, CNV-espresso achieved a significant effect on improving the precision for small CNVs with less than 10 targets, which just fills the current gap in accurately identifying small CNVs from exome sequencing data. Meanwhile, CNV-espresso can significantly exclude up to 71% of false-positive CNVs while retaining the majority of true positive calls (**Figure 5 D, E, F**). We applied CNV-espresso directly to the raw CNV calls and achieved similar filtering results as the traditional quality score filtering approach. Furthermore, by combining CNV-espresso and the traditional filtering approach together, we can exclude additional false-positive calls which were missed by the traditional filtering approach and achieve a higher precision result.

**Figure 5.**
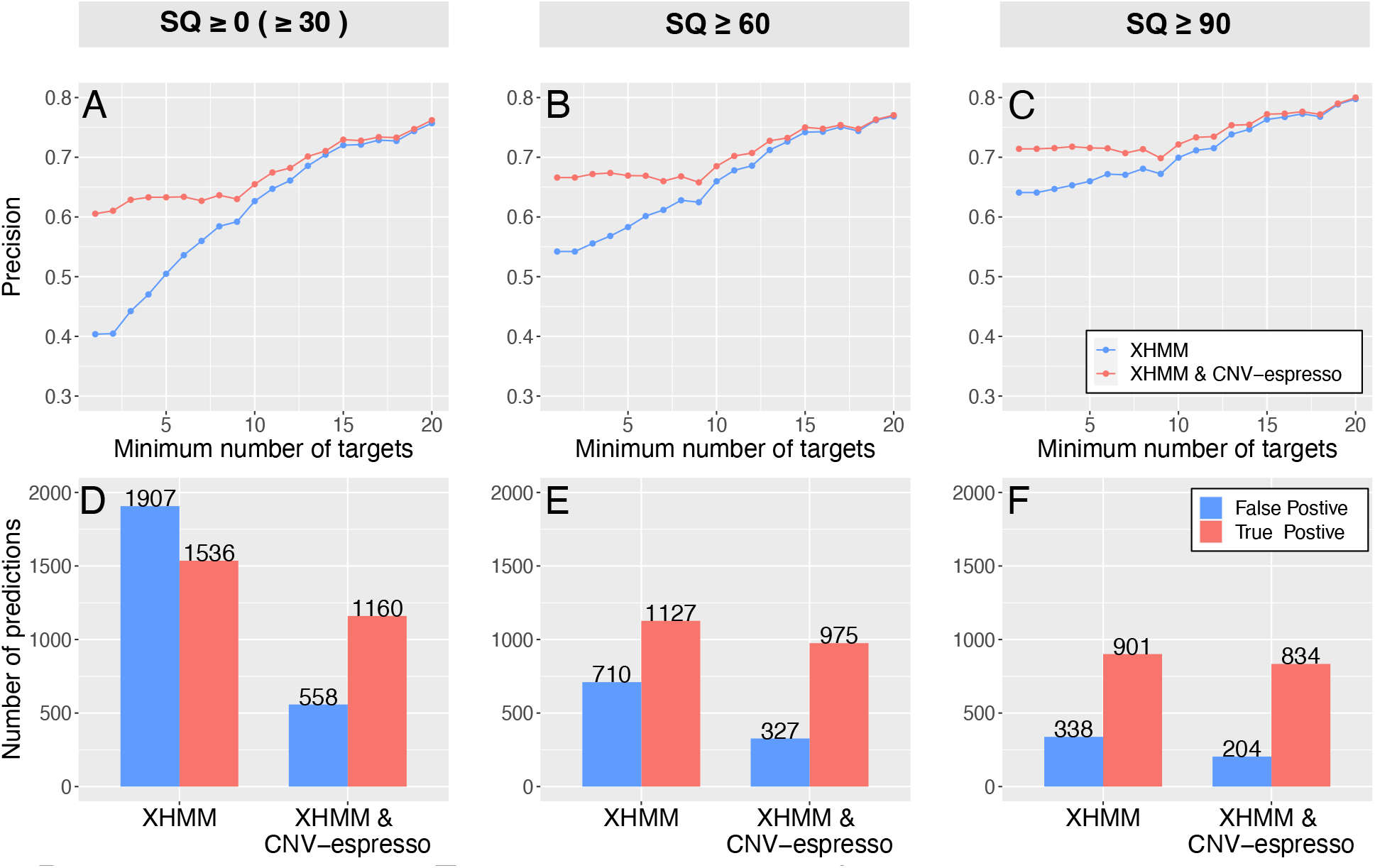
Changes in precision and the number of CNV predictions before and after *in silico* confirmation by CNV-espresso. This analysis was conducted with different benchmark CNV call sets: (A, D) raw CNV predictions from XHMM. Note that all SQ scores of the raw CNV predictions are greater than 30 in our initial call set; (B, E) CNV predictions with SQ scores greater than or equal to 60, which is a recommended score for filtering in the XHMM pipeline; (C, F) CNV predictions filtered by a stringent score, SQ ≥ 90. For each benchmark call set, we checked the precision changes with different CNV sizes, as well as the number of true positive and false positive predictions before and after *in silico* confirmation by CNV-espresso.

### Results on experimentally validated dataset

To assess our model’s compatibility under different capture kits and experimental batches, we applied CNV-espresso to *in silico* confirm CNVs on a real experimentally validated dataset from a congenital heart disease study (38). We obtained 24 true positive CNVs validated by digital droplet polymerase chain reaction (ddPCR). The corresponding exome sequencing samples of these 24 CNVs were captured by Nimblegen SeqCap Exome V2 chemistry and sequenced on the Illumina HiSeq 2000 platform. Sequence reads were aligned to the human reference genome hg19 as described (39). We used our trained model to validate the CNV predictions for these 24 true CNVs. We found CNV-espresso can successfully confirm 23 of the 24 CNVs previously picked up by exome sequencing technology with a true positive rate of 96%. Among them, for the CNVs with more than 15 capture targets, the true positive rate reached 100%. Overall, results showed that our model can achieve a good true positive rate on the experimentally validated dataset.

## Discussion

In this study, we present a new deep transfer learning method, CNV-espresso, for *in silico* confirming CNV predictions from exome sequencing data. In genomic studies using exome sequencing data, an indispensable step in CNV analysis is to manually visualize the images of putative CNVs that contain visual information about read depth. The core idea of CNV-espresso is to use deep learning models optimized for image recognition to capture implicit logics in manual visualization by humans. For each CNV candidate predicated by exome-sequencing-based CNV callers, CNV-espresso encodes normalized read depth signal for the sample of interest and selected reference samples into an image. Then, CNV-espresso adopts transfer learning with a pre-trained CNN model and fine-tuning the model by a large-scale trio-exomes dataset. We evaluate the performance of CNV-espresso on an independent dataset with both exome sequencing and whole genome sequencing data, and an experimentally validated dataset. Our results show that CNV-espresso can achieve a high performance for filtering out false positives without substantial loss of sensitivity. It can perform robustly on confirming both of deletion and duplication. Furthermore, CNV-espresso can successfully *in silico* confirm CNV predictions among different size categories. It can precisely exclude up to 71% of false-positive calls while retaining most true positive calls. Importantly, CNV-espresso can successfully confirm small CNVs with only a few targets, which is currently one of the biggest challenges in calling CNVs with exome area. Finally, our results also show that CNV-espresso can work compatibly with different capture kits and experimental batches and achieve a high true positive rate with ddPCR experimental results.

There are several limitations to our study. First, the main signal used by CNV-espresso is the depth contrast between sample of interest and a selected set of reference samples. The presence of the same CNV in the reference samples would decrease the signal. Therefore, CNV-espresso is optimized for confirming rare CNVs. Second, CNV-espresso is designed to confirm CNV calls made by other methods. It can reduce false positives without substantial loss of sensitivity, and by design it cannot improve sensitivity.

## Methods

### Samples and CNV calling

We obtain exome sequencing data from 27,270 family-based samples in the autism spectrum disorder cohort. The samples are processed with custom NEB/Kapa reagents, the IDT xGen capture platform, and sequenced on the Illumina NovaSeq 6000 system by Regeneron. Sequence reads are aligned to the human reference genome hg38.

We identify CNVs from exome sequencing data by using XHMM (13), CANOES (16) and CLAMMS (19), three CNV callers that have different statistical models. We use principal component analysis to illustrate samples and check whether the read depth signals of these samples have been significantly affected by batch effects. After passing the batch effects inspection, we randomly split samples into ten groups (for XHMM and CANOES) or two groups (for CLAMMS) according to the computational complexity of different CNV callers. Note that in the random grouping process, we require members in the same trio to be assigned into the same group for CNV calling. In this study, we use all default parameters in CANOES and CLAMMS. For XHMM, we appropriately adjust the ‘--maxTargetSize’ parameter to keep 12 capture target probes longer than 10,000 bp not being excluded as outliers. Except that, we keep all other parameters by defaults.

In addition to the 27,270 exome sequencing samples mentioned above, we also obtained additional 569 individuals with both exome sequencing and whole genome sequencing data from the SPARK cohort (12). For these 569 individuals, we used XHMM (13) to call putative CNVs from exome sequencing data; we used Canvas (36) and Manta (37) to identify CNVs from whole genome sequencing data with default software parameters. We filtered the raw CNV calls from Canvas and Manta by recommended quality control criteria. Specifically, we excluded CNVs with quality scores smaller than 7 in Canvas’ results; we only kept deletions and duplications located on autosomal and sex chromosomes (excluded any predicted variants on contigs) in Manta’s results. In addition, we required ‘PASS’ in the ‘FT’ field of the VCF file, which indicates that all the filters of Manta have passed for the corresponding sample.

### Read depth calculation and GC content normalization

To avoid the additional variance caused by the extremely long (≥1000bp) target regions, we first divide those extremely long targets into serval equal size 500∼1000 bp windows (referred to as “target” in the manuscript). This approach can also benefit to confirm CNVs located at part of the long target regions (19). We then employ Mosdepth (40) to calculate the read depth signal for each target of each sample. Users can also use other read depth generators to perform calculations or even directly take the read depth files from CLAMMS as input. It is well known that the real CNV signals in the exome sequencing data are significantly affected by noises. The noise signals include GC content, sample mean coverage, target size, target mean coverage, CNV frequency in population and batch effects etc. (13). These factors can affect read depth signal globally or locally. Among them, GC content and sample mean coverage are known to affect the read depth signal globally. Thus, we normalize the GC content and sample overall coverage for each sample by a median approach as:

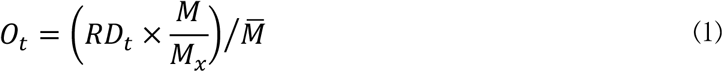

where *RD*_*t*_ is to the raw read depth at the *t* th exon. *M*_*x*_ is the median read depth value of all exons with the same GC content *x* as the exon, and *M* is the overall median read depth of all exons in a sample, while 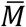 is the overall mean read depth of all exons in a sample. *O*_*t*_ is the normalized read depth value of the exon. **Figure 6** shows the read depth signal before and after GC content normalization.

**Figure 6.**
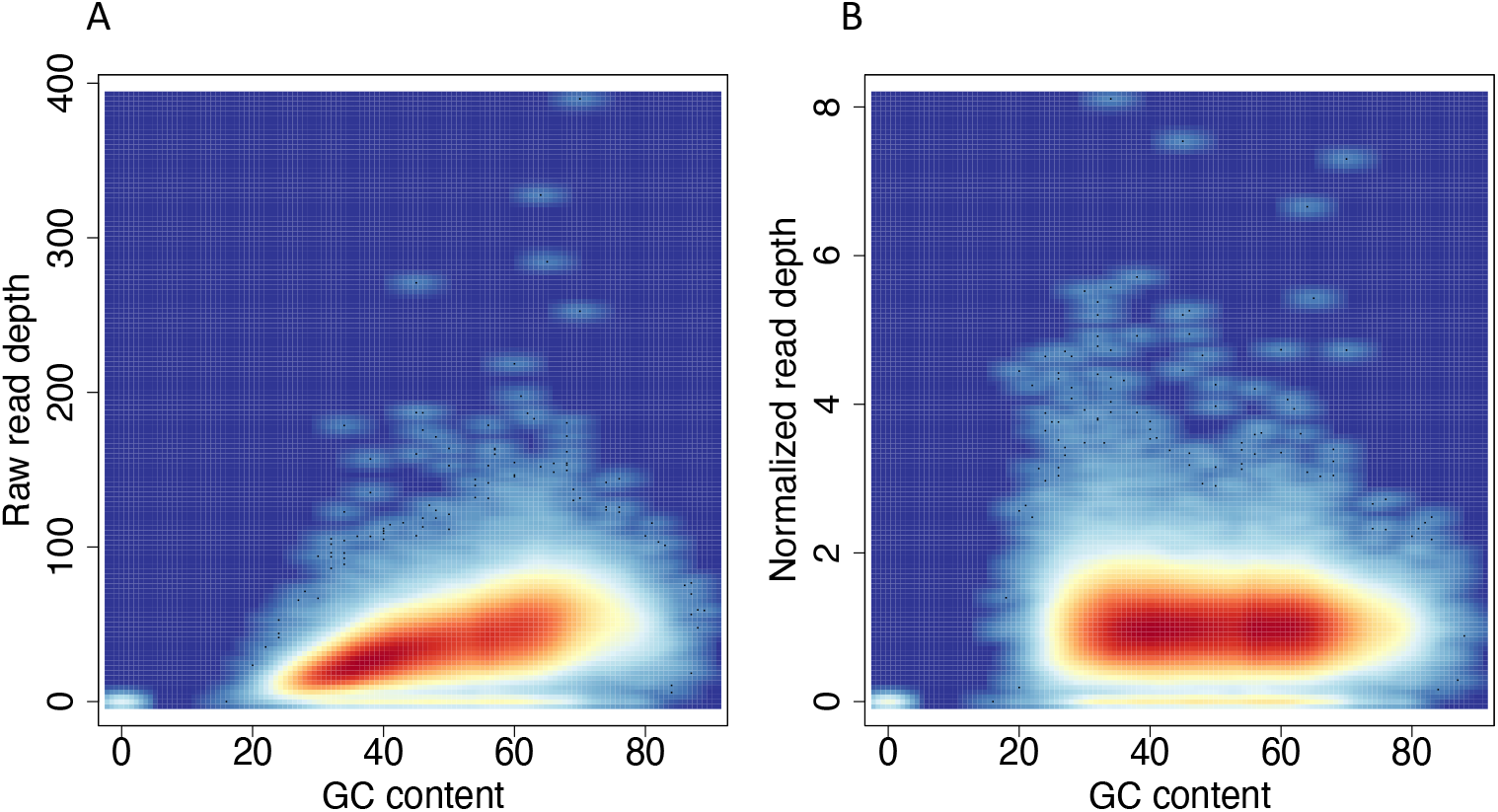
GC normalization. (A) Correlation between GC content and the raw read depth before normalization. (B) Correlation between GC content and normalized read depth.

### Reference samples selection

For other known and unknown factors locally affecting read depth signals, we assume that these factors contribute equally on the same batch of samples in the given genomic regions. Especially for those reference samples with high similarity to the case sample. The rare CNV signals can be inferred from the contrast relationship between the case and reference samples on each target of the exome. For instance, previous studies successfully detected rare CNVs by assuming the normalized number of read counts or read depth values in exome target regions follow a negative binomial or similar distributions (15,16,21). Inspired by these methods, we calculate the correlation coefficients of the case with other samples and select the 100 (by default) highest pairwise correlation samples as reference samples.

### Image encoding

For each CNV candidate predicted by CNV callers, we encode the read depth signals of the case and its corresponding reference samples into an image. The X-axis of the image refers to the CNV coordinate in the human genome, and the Y-axis of the image refers to the normalized read depth value. To avoid image abnormalities caused by the fluctuations in distance changes between adjacent targets, we take the logarithmic transformation for the differences between two adjacent targets on the X-axis and the normalized read depth values of all exons on the Y-axis as equations (2) and (3) respectively.

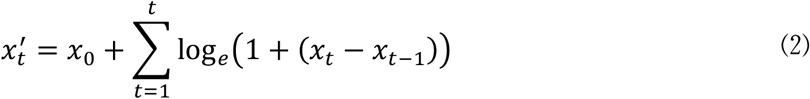

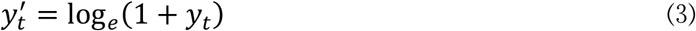

where *x*_*t*_ is the genomic coordinate at the middle of *t* th exon, and *y*_*t*_ refers to the normalized read depth value at *t* th exon. To avoid the undefined error, we use the natural logarithm of one plus the input in the transformation.

In the image, each dot refers to a target region, and we connect every two adjacent dots of the sample with a straight line. We use RGB color to distinguish different type of samples. Here we use blue for the case sample and gray for reference samples. **Figure 7** illustrates example images of deletion and duplication after encoding. Thus, we convert the CNV confirmation task into an image classification question.

**Figure 7.**
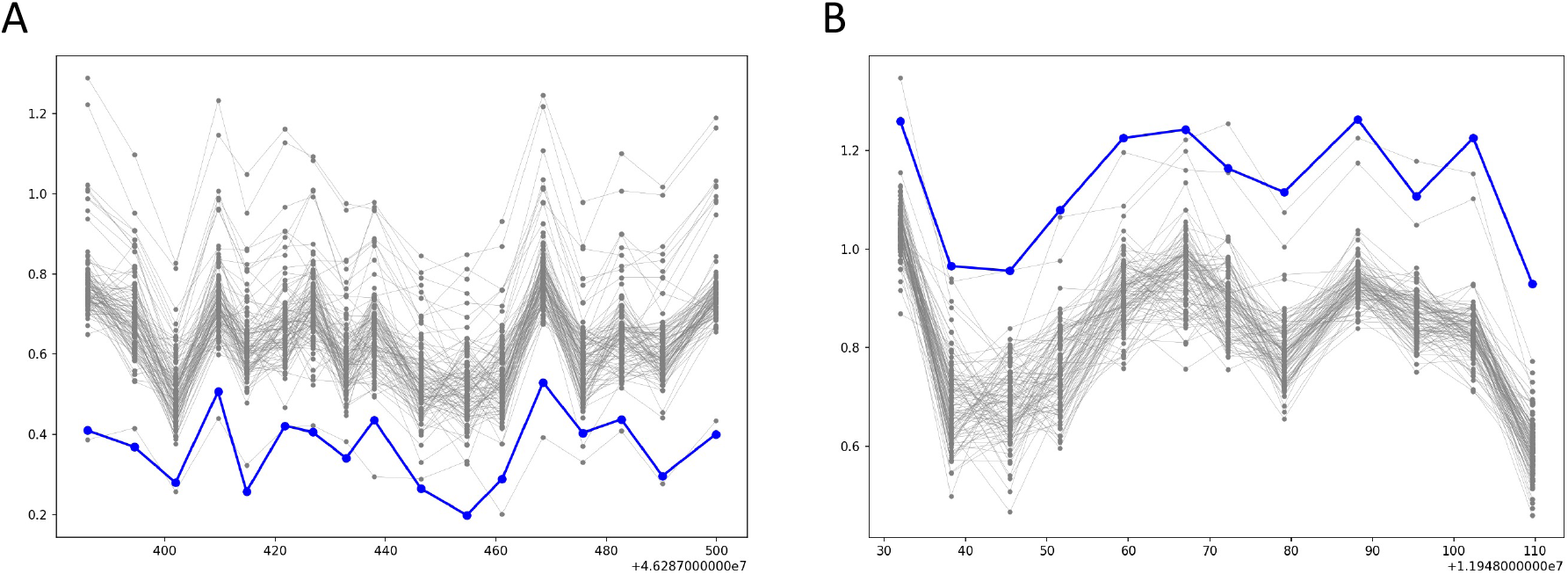
Representative images for CNV predictions. (A) Deletion. (B) Duplication.

### Transfer learning

In this study, we leverage the transfer learning and fine-tuning strategy to *in silico* confirm rare CNV calls from exome sequencing data. As requested by the pre-trained base model, we resize our input CNV images to 224×224 pixels. Given that the purpose of our study is to train a three-class (deletion, duplication, and diploid) deep learning classifier, we exclude the top layer of MobileNet v1 as the base model, then add a global average pooling layer and a dense layer with Softmax activation function.

We apply the typical transfer learning and fine-tuning procedures from Keras (41). Specifically, we first take layers from the pre-trained MobileNet v1 base model and freeze all the weights as non-trainable to avoid destroying any of the information they already contained. Then, we train the weights of newly added layers using our labeled dataset. Once the model converges, we set all the weights of the model trainable and retrain the whole model end-to-end with a very low learning rate (10e-5) by Adam optimizer (42). We train the CNN model in batches of 32 images for 20 epochs on a GPU server [GeForce GTX 1080 GPU, 8119MiB RAM]. Early stopping was set by monitoring the value of ‘loss’ and three epochs with no improvement after which training will be stopped. We select the model with the highest accuracy on the testing data as the final model.

### Evaluation metrics

We evaluate the performance of the refined model on the testing data. Specifically, we use the refined model to predict the probabilities of the three copy number states (deletion, duplication, and diploid). We select the state corresponding to the maximum probability value as the predicted label. We treat a CNV in the testing set as true positive (TP) if its predicated label matches the corresponding label. We count any true CNVs without a matched predicted label as false negatives (FN), while any predicted CNVs without a matched true label as false positives (FP). We evaluate the performance of CNV-espresso by using F1-score and area under the curve (AUC). We calculate the F1-score as:

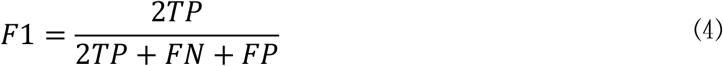

## Data availability

The software is implemented in python and freely available from the GitHub repository at https://github.com/ShenLab/CNV-Espresso.

## Acknowledgments

We thank Dr. Haicang Zhang, Joseph U. Obiajulu, Dr. Chang Shu, Dr. Bingsong Zhang, and Jiayao Wang for helpful discussions. The work was partly supported by NIH grants R01GM120609 and R03HL147197 and Simons Foundation (SIMONS606450).

The exome and whole genome sequencing data sets from the SPARK project used in this work are available at https://base.sfari.org (application required). The SPARK initiative is funded by the Simons Foundation as part of SFARI. We are extremely grateful to the thousands of individuals and families who are participating in SPARK. We wish to thank the sites, staff, and volunteers of the SPARK Clinical Site Network and SFARI for their invaluable contributions.

