## Supplementary figures and table for "Accurate *in silico* confirmation of rare copy number variant calls from exome sequencing data using transfer learning"

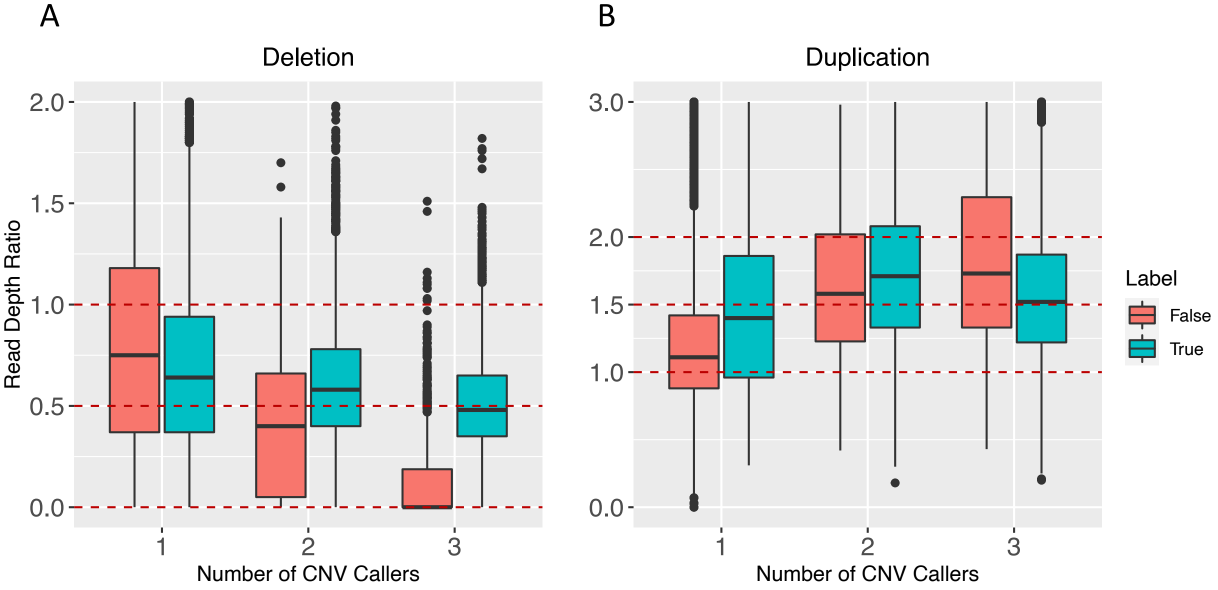


**Supplementary figure 1. Box plots illustrating the distribution of read depth ratio between CNV coordinates and their boundary regions versus different number of CNV callers in the labeled dataset.** We illustrated those CNV predictions by deletion (A) and duplication (B) separately and classified them by "true" or "false" label in different colors. For each CNV prediction, we first counted the number of exome targets located within the CNV coordinates, then selected half of this number of targets from each side of boundary regions. The read depth ratio was calculated as the average read depth within the CNV coordinates divided by the average read depth in left and right boundary regions. In theory, the read depth ratio should be close to 0 for homozygous deletions and 0.5 for heterozygous deletions; 1 for diploid regions, 1.5 and more for duplications. We observed that in the "false" label set, as the deletions (duplications) were concordantly identified by more CNV callers, the read depth ratio has an obviously decreasing (increasing) trend. This means at least partly of CNV predictions concordantly identified by multiple CNV callers in the "false" label set are likely to be real, which need to be excluded from the final "false" label set.

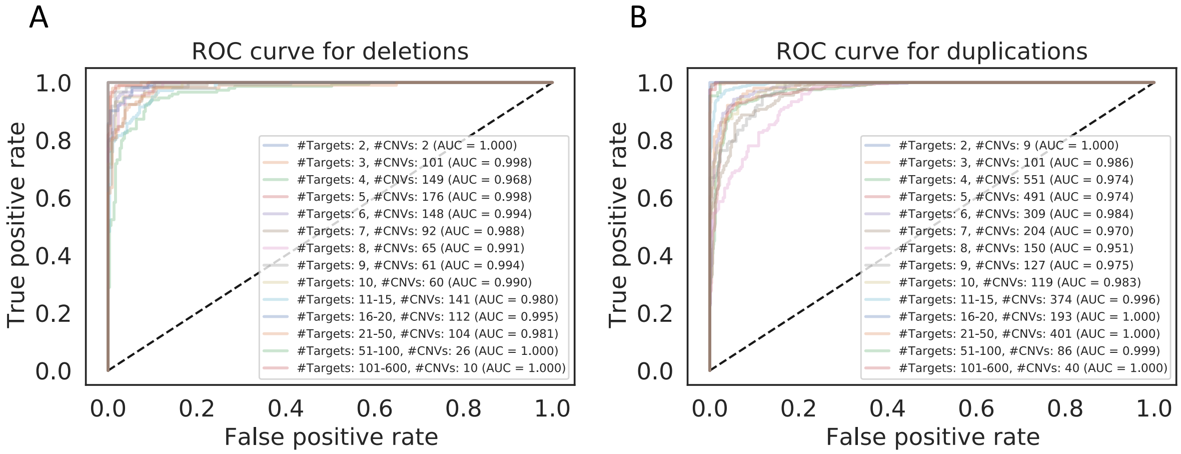


**Supplementary figure 2. Performance comparisons on different CNV types and number of targets.** (A) ROC curves for deletions with different number of targets. (B) ROC curves for duplications with different number of targets.

**Supplementary table 1. Filtering criteria for generating training data using CNV calls made by three different methods**

| Label in training data | XHMM | CANOES | CLAMMS |
|---|---|---|---|
| True (inherited calls) | Offspring.SQ ≥ 60 & One.Parent.NQ ≥ 60 & Other.Parent.SQ ≥ 60 | Offspring.SQ ≥ 70 & One.Parent.NQ ≥ 70 & Other.Parent.SQ ≥ 70 | EQ > 0 & CNVs were shared (≥ 1 bp) by other CNVs in the parents |
| False (Mendelian errors) | Offspring.SQ ≥ 10 & Parents.NQ ≥ 60 | Offspring.SQ ≥ 10 & Parents.NQ ≥ 70 | EQ > 0 & SQ ≤ 80 & CNVs were NOT shared (≥ 1 bp) by any CNVs in the parents |
| Additional filtering criteria | 1. Not located in SD regions, and<br>2. Not located in low Mappability (<0.25) regions, and<br>3. Not located in extreme GC content (<0.3 or >0.7) regions, and<br>4. Exclude CNVs identified by two or three callers in the dataset with 'False' label | | |

SQ is the statistical quality score which refers to the Phred-scaled probability of some CNVs in the interval; NQ is the Phred-scaled probability of not being CNV in the interval, and EQ is the Phred-scaled probability of the exact CNV event in the given interval.
